# T-ALL Proliferation Relies on Glutamine Uptake and EAAT1-dependent Conversion of Glutamine to Aspartate and Nucleotides

**DOI:** 10.1101/2020.02.08.939694

**Authors:** Vesna S. Stanulović, Shorog Al Omair, Michelle A.C. Reed, Sandeep Potluri, Jennie Roberts, Tracey A. Perry, Sovan Sarkar, Guy Pratt, Ulrich L. Günther, Christian Ludwig, Maarten Hoogenkamp

## Abstract

T-cell acute lymphoblastic leukaemia (T-ALL) is a cancer of the immune system. Approximately 20% of paediatric and 50% of adult T-ALL patients have refractory disease or relapse and die from the disease. To improve patient outcome new therapeutics are needed. With the aim to identify new therapeutic targets, we analysed the metabolic adaptations that T-ALL cells exhibit and found that glutamine uptake is essential for their proliferation. Isotope tracing experiments showed that glutamine fuels aspartate synthesis through the TCA cycle and that glutamine and glutamine-derived aspartate together supply three nitrogen atoms in purines and all but one atom in pyrimidine rings. We show that the glutamate-aspartate transporter EAAT1, which is normally only expressed in the CNS, is crucial for glutamine conversion to nucleotides and that T-ALL cell proliferation depends on EAAT1 function. Through this work, we identify EAAT1 as a novel therapeutic target for T-ALL treatment.

## Introduction

T-cell acute lymphoblastic leukaemia (T-ALL) are haematological malignancies of the T-cell lineage. Subtypes are grouped based on immunophenotype, or more recently, gene expression profiling that correlates with the underlying genomic aberrations^1,2^. T-ALL occurs both in adults and children and, despite improved treatment outcome in the last decades, up to 20% of paediatric and 50% of adult T-ALL patients have refractory disease or relapse^3-6^. Alternative treatment options are not available for these patients and, therefore, novel therapeutic targets are needed.

For a cancer to proliferate, cells need to increase their size and replicate their DNA, which requires large quantities of proteins, lipids and nucleotides, as well as energy. Cell proliferation and survival require metabolic adaptations, involving glycolysis, glutaminolysis and the PI3K-AKT-mTOR pathway^7-12^. The compilation of metabolic changes results in a distinct metabolic phenotype and determination of the rate-limiting steps that support the oncogenic metabolic adaptations can lead to the identification of novel therapeutic targets.

With the aim to identify common oncogenic adaptations, we investigated T-ALL metabolism, using four cell lines, representing different stages of T-cell development, and primary patient samples. We found that T-ALL cells uptake glutamine and use it to generate aspartate. Normally, glutamine and aspartate are used as substrates for nucleotide production but in T-ALL glutamine alone provides all of the nitrogen atoms and all but one carbon atom in the pyrimidine ring, and three of the four nitrogens in purines. Furthermore, we determined that T-ALL cells express the glutamate-aspartate antiporter EAAT1 (encoded by *SLC1A3*), which is present in 95% of T-ALL patients^2^. In the healthy adult body, EAAT1 is only present in the neurons and glia of the central nervous system where it removes cytotoxic glutamate from the glutamatergic synapses in exchange for aspartate^13,14^. In T-ALL, EAAT1 is required for glutamate import into mitochondria in exchange for glutamine-derived aspartate which is then used as a substrate for nucleotide production in the cytoplasm. We show that T-ALL survival depends on glutamine uptake and EAAT1 function, which validates EAAT1 as a therapeutic target for treating T-ALL.

## Experimental Procedures

### Cell Culture

T-ALL cell lines were grown in RPMI 1640medium (Merck) supplemented with 10% FCS, 5U/ml Pen/Strep, 2mM Glutamine or GlutaMax (LifeTechnologies) and 0.075mM 1-Thyoglycerol (Merck) in a humidified incubator at 37°C and 5% CO_2._ Where indicated, UCPH-102 (Abcam) was added to a final concentration of 25μM. For labelling experiments, glutamine-free RPMI was supplemented with 2mM [1,5-^15^N]-L-Glutamine or [3-^13^C]-L-Glutamine (Merck).

*In vitro* differentiation of human T-cell progenitors was performed from CD34^+^ haematopoietic stem/progenitor cells. Mononuclear cells were isolated from cord blood using Lymphoprep™ (StemCell Technologies). Red blood cells were lysed using 1x Lysing buffer (BD Biosciences) at RT for 10 minutes. CD34^+^ cells were isolated using a human CD34 MicroBead Kit (Miltenyi Biotec), according to the manufacturer’s protocol using LS columns (Miltenyi Biotec). CD34^+^ cells were co-cultured on 80% confluent OP9-DL4 cells in αMEM medium, supplemented with 5 ng/ml human FLT3L (Peprotech) and 5 ng/ml human IL7 (Peprotech). Cells from the *in vitro* human T-cell differentiation were sorted at specific time points for RNA isolation and subsequent RNA sequencing. Cells with the CD7^+^ CD5^-^ CD1a^-^ immunophenotype were sorted at day 7, CD7; CD5^+^ CD1a^-^ at day 14, and CD7^+^ CD5^+^ CD1a^+^ CD3^-^ at day 21. Purification of cell populations was performed using a BD FACSAria Fusion (BD Biosciences) cell sorter.

### Expression analysis

RNA isolation and cDNA synthesis was performed as previously described (Stanulovic et al 2017). Quantitative PCR (qPCR) was performed using an ABI 7500 Real-Time PCR System. Quantitation was carried out using a standard curve of serial dilutions and relative to rDNA. Primers used for quantification:SLC1A3F 5’AAGAGAACAATGGCGTGGAC,SLC1A3R 5’ATTCCAGCTGCCCCAATACT, 18SrRNAF 5’GGCCCGAAGCGTTTACTTTGA, 18SrRNAR 5’GAACCGCGGTCCTATTCCATTA.

### RNASeq library preparation

RNASeq libraries were prepared and indexed using the TruSeq Stranded mRNA Sample Preparation Kit LH (Illumina) according to the manufacturer’s protocol. Quality control of the libraries was carried out by Agilent Bioanalyzer using the High Sensitivity DNA analysis kit, and quantitation of the libraries was carried out using the Kapa Biosystems Illumina library Quantitation kit. Libraries were pooled and sequenced as 100 nt paired-end on Illumina HiSeq 2500 sequencer at a depth of approximately 30 million reads per library.

### RNA-seq data analysis

Reads acquired from RNASeq were mapped as stranded libraries to the human genome hg38 (GRCh38), using HISAT2 using the public server at usegalaxy.org^15,16^. Transcripts were assembled using StringTie and GENCODE gene annotation with quartile normalisation and effective length correction^17^. Assembled transcripts were merged and normalised before differential gene expression was determined by DESeq2. Duplicate biological replicates were used. Gene differential expression, between T-ALL samples and in vitro differentiated samples, and FPKM gene expression files were used to select the gene IDs that were significantly differential expressed. Pearson complete linkage hierarchical clustering and heat maps were computed by Multi Experiment Viewer software^18^. Gene ontology enrichment was performed using DAVID 6.8 beta^19,20^. GO terms for biological processes with Modified Fisher Extract P-value <0.05 were considered significant, categories with redundant terms were filtered out, and the top 10 are presented. To visualize the distribution of transcripts across the reference genome, the DNA strand specific bam files were transformed to bigwig files and uploaded as composite tracks to the UCSC browser. Data has been deposited at NCBI-GEO GSE101566.

### Western blot analysis

Cell extracts were prepared by lysing 10^7^ cells in 50µl cell lysis buffer (20mM TRIS pH8, 150mM NaCl, 2mM EDTA, 0.5% NP40) or RIPA buffer (50mM TRIS pH8, 150mM NaCl, 0.5% deoxycholic acid, 1% NP40, 0.1% SDS) on ice for 20 min. The insoluble fraction was precipitated by centrifuging at 20,000 g for 10min. Protease inhibitor and PhosSTOP (Roche) were used 1:100.

Proteins were separated on a 4-12% gradient Bis-Tris Plus Bolt Mini Gels (LifeTechnologies)and transferred to nitrocellulose membranesGels were stained with PonceauS to confirm equal loading prior to o/n incubation with antibodies. Primary antibodies raised against EAAT1 (D4166; Cell Signalling) and GFP (GF28R; eBiosciences) were used at a final concentration of 1µg/ml. Secondary antibodies, IRDye 680RD or 800RD (Li-Cor), were used at a 0.5µg/ml. Westerns were visualised using an Odyssey CLx Imager (Li-Cor).

### Immunofluorescent Staining

Cells were fixed with 2% formaldehyde for 10min and washed twice in PBS/0.05%Tween/2%FCS, followed by o/n incubation with 1μg/ml primary antibody. Cells were washed and incubated with 1µg/ml secondary antibody conjugated to Alexa dyes (LifeTechnologies). Immuno-stained cells were deposited onto the glass slides using a Cytospin III centrifuge (Shandon). MitoTracker Red CMXRos (Invitrogen) was used for mitochondria staining as per manufacturer’s instruction. Slides were dried, covered with Prolong Gold Anti-Fade DAPI reagent (LifeTechnologies) and imaged using a Zeiss LSM880 Confocal microscope.

### SLC1A3 knockdown

SLC1A3 was amplified by PCR from pcDNA3-EAAT1. The PCR product was cloned into MigR1 in front of IRES-GFP^21^. Short hairpin sequences targeting *SLC1A3* were designed using http://cancan.cshl.edu/RNAi_central/RNAi.cgi?type=shRNA^22^. Designed shRNA (shSLC1A3_1 ACCATATCAACTGATTGCACAG, shSLC1A3_2 GGGTAACTCAGTGATTGAAGAG, shSLC1A3_3 GTGGCACACAATCCTATAAATG, shSLC1A3_4 AGGCCTCAGTGTCCTCATCTAT and shSLC1A3_5 CACTCCTCAACTGATGATAGAC) were embedded into mir30 and cloned into pMSCVhygro^23^. The mouse fibroblast cell line PlatE was transfected using *Trans*IT-LT1 (Mirusbio) with MigRI-SLC1A3 and MSCVhyg_shSLC1A3, MSCVhyg_shGFP, or MSCVhyg_shFF3^22,24^.

Functional shRNA was cloned into piggybac transposon inducible expression vector PB_tet-on_Apple_shGFP using *Hind*III and *Kpn*2L (ThermoFisher)^25^. T-ALL cell lines were electroporated with pB_shSLC1A3 and p*CAGG-PBase* ^26^, expanded and selected with puromycin (ThermoFisher). Expression of shRNA was induced with 0.1μg/ml doxycycline (Merck). Cells were counted every other day and the cell concentration was adjusted to 0.4×10^6^/ml.

### Patient Samples

The patients’ cells used in this study were from diagnostic samples from presentation cases before treatment. They were obtained from the Queen Elizabeth Hospital Birmingham, Birmingham, UK with ethical approval from the NHS National Research Ethics Committee (Reg:10/H1206/58). Cytogenetic abnormalities and sample immunophenotype were determined at the time of disease diagnosis at the West Midlands Regional Genetics Laboratory, Birmingham Women’s NHS Foundation Trust, Birmingham, UK.

Mononuclear cells were purified from peripheral blood by differential centrifugation using Lymphoprep (Axis-Shield UK). Undifferentiated blast cells were isolated using anti-human CD34 (T-ALL_1 and _2) or CD7 (T-ALL_3) MACS microbeads (Miltenyi). For T-ALL_2, CD34^+^ cells were further sorted by FACS for CD7^+^ using anti-human CD7-FITC antibody (CD6-6B7; BioLegend).

### Intracellular NMR spectroscopy

Methanol/chloroform extraction was used to prepare polar extracts from 5×10^7^ cells. Cells were washed twice with PBS, pelleted by centrifugation and resuspended in ice cold methanol. After adding water and chloroform, samples were vortexed and incubated on ice for 10min. Following centrifugation, the polar phase was dried in a vacuum concentrator. Pellets were resuspended in 50µl NMR buffer (100 mM sodium phosphate, 500µM Sodium 3-(trimethylsilyl)propionate (TMSP; Merck), 10% D_2_O, pH7.0)and and transferred to 1.7mm NMR tubes. NOESY 1D spectra with water pre-saturation were acquired using the standard Bruker pulse sequence noesygppr1d on a Bruker 600MHz spectrometer with a TCI 1.7mm z-PFG CryoProbe™. The sample temperature was set to 300K. The ^1^H carrier was on the water frequency and the ^1^H 90° pulse was calibrated at a power of 0.326W. Key parameters were: spectral width 12.15ppm/7288.6Hz; complex data points, 16384; interscan relaxation delay, 4s; acquisition time, 2.25s; short NOE mixing time, 10ms; number of transients, 256; steady state scans, 4.

### Growth-media metabolite uptake and release

Cells were grown under normal conditions. Growth medium was collected and supplemented with 10% D_2_O and 1mM TMSP. Samples were transferred to 5mm glass NMR tubes and spectra were acquired at 300K, using a Bruker 500 MHz spectrometer equipped with a TXI probe. The ^1^H carrier was on the water frequency and the ^1^H 90° pulse was calibrated at a power of 12.9W. Measurements were carried out after locking on deuterium frequency and shimming. The standard Bruker 1D NOESY pulse sequence (noesygppr1d) with water saturation was used. Spectra were acquired with 64 transients and 4 steady state scans.

### Live-cell real-time NMR spectroscopy

Cells were resuspended at 10^6^/ml growth media containing 0.1% low melting agarose (Sigma) with 1mM TMSP and 10% D_2_O. Samples were loaded into 5mm NMR tubes and measurements were collected every 8.4 minutes, for a total of 100 time points. CPMG (Carr-Purcell-Meiboom-Gill) 1D spectra with water pre-saturation were acquired using the standard Bruker pulse sequence cpmgpr1d on a Bruker 500 MHz spectrometer with a TXI ^1^H/D-^13^C/^15^N probe at 310K. The ^1^H carrier was on the water frequency and the ^1^H 90° pulse was calibrated at a power of 12.9W. For the CPMG T_2_ filter, a T_2_ filter time of 68 ms arose from 100 loops over a 680 µs echo time between 180 pulses. Other key parameters were: spectral width 12.02 ppm/6009.6 Hz; complex data points, 16384; interscan relaxation delay, 4s; acquisition time, 2.73s; short NOE mixing time, 10ms; number of transients, 64; steady state scans, 4.

### NMR spectroscopy data processing and analysis

NMR spectroscopy data was processed in Topspin (Bruker Ltd, UK) and MetaboLab (version 20910688) in MATLAB (version R2015b)^27^. All spectra were aligned to TMSP, a spline baseline correction was applied, the water and TMSP regions were excluded, and the total spectral area (TSA) of each spectrum was scaled to 1. To compare metabolite concentrations, a well-resolved peak was picked for each metabolite in the first spectrum and peaks were picked in the other spectra in an automated manner using in-house subroutines of MetaboLab (version 20910688). Spectral assignments were made using Chenomx software and acquired chemical standards.

### Mouse studies

Animal experiments were performed at the University of Birmingham Biomedical Services Unit under an animal project licence (PP8841933) in accordance with UK legislation. Female NOD.Cg-Prkdcscid Il2rgtm1Wjl/SzJ (NSG) mice aged 8-9 weeks at study commencement were used for the xenograft model. CCRF-CEM cells were transfected with pBshSLC1A3_2 or pBshGFP control vector and pCAGG-PBase as described above. After selection, 3×10^5^ cells per mouse were injected *intra venous* using the tail vein. When approximately 1% engraftment was observed, the next day all animals were transferred to a diet supplemented with 0.625g/kg doxycycline hyclate.

## Results

### T-ALL metabolic phenotype

Based on the prevalence, the risk of refraction/relapse, and the stage at which T-cell development was blocked, we chose one cell line that immuno-phenotypically belongs to the ETP T-ALL group (ARR) and three other cell lines that are blocked at different stages of T-cell development (DU.528, HSB2 and CCRF-CEM)^28^. To assess leukaemic adaptation, we compared the gene expression of these cell lines to the gene expression of *in vitro* differentiated T-cell progenitors at the CD7^+^ CD5^-^ CD1a^-^, the CD7^+^ CD5^+^ CD1a^-^, and the CD7^+^ CD5^+^ CD1a^+^ CD3^-^ stage of differentiation. Hierarchical clustering of the differentially expressed genes identified 7 clusters (Figure 1A). Genes suppressed in the T-ALL samples were grouped in cluster 1 (C1) and C5. Gene functional annotation analyses indicated several terms related to mitochondrial function (Figure 1B). This is in line with the observation that T-ALL cells exhibit depletion of ATP and chronic metabolic stress^29^.

**Figure 1.**
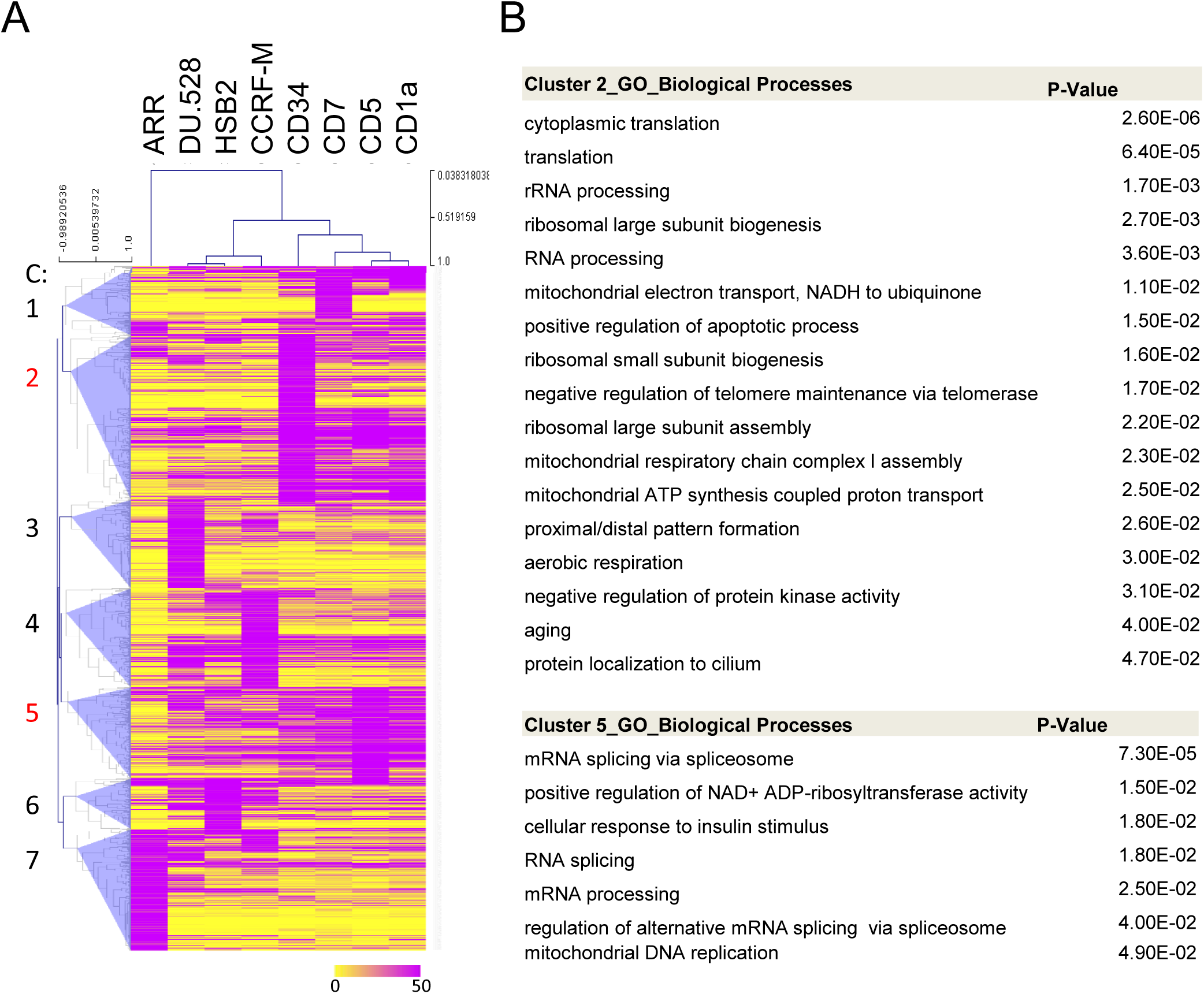
Differential gene expression and gene ontology analysis for T-ALL cell lines. **A)** Heat map showing hierarchical clustering of RNAseq gene expression data, based on Pearson correlation with complete linkage clustering of all differentially expressed genes. Clusters are marked by blue triangles and numbered from 1 to 7. Scale bar represents log2 FPKM values. **B)** Gene ontology enrichment analysis for clusters 2 and 5. Terms are ordered based on Modified Fisher Extract P-value.

We therefore investigated the metabolism of the TALL cell lines. Metabolite uptake was assessed by measuring metabolite levels in the media, 24h after medium change, by NMR spectroscopy and comparing it to the levels present in the media alone. We identified 22 metabolites with significantly changing levels in at least one of the T-ALL cell lines and found that all four cell lines utilised lysine, glucose, phenylalanine and glutamine and released lactate, pyruvate, glutamate and pyroglutamate (Figure 2A).

**Figure 2.**
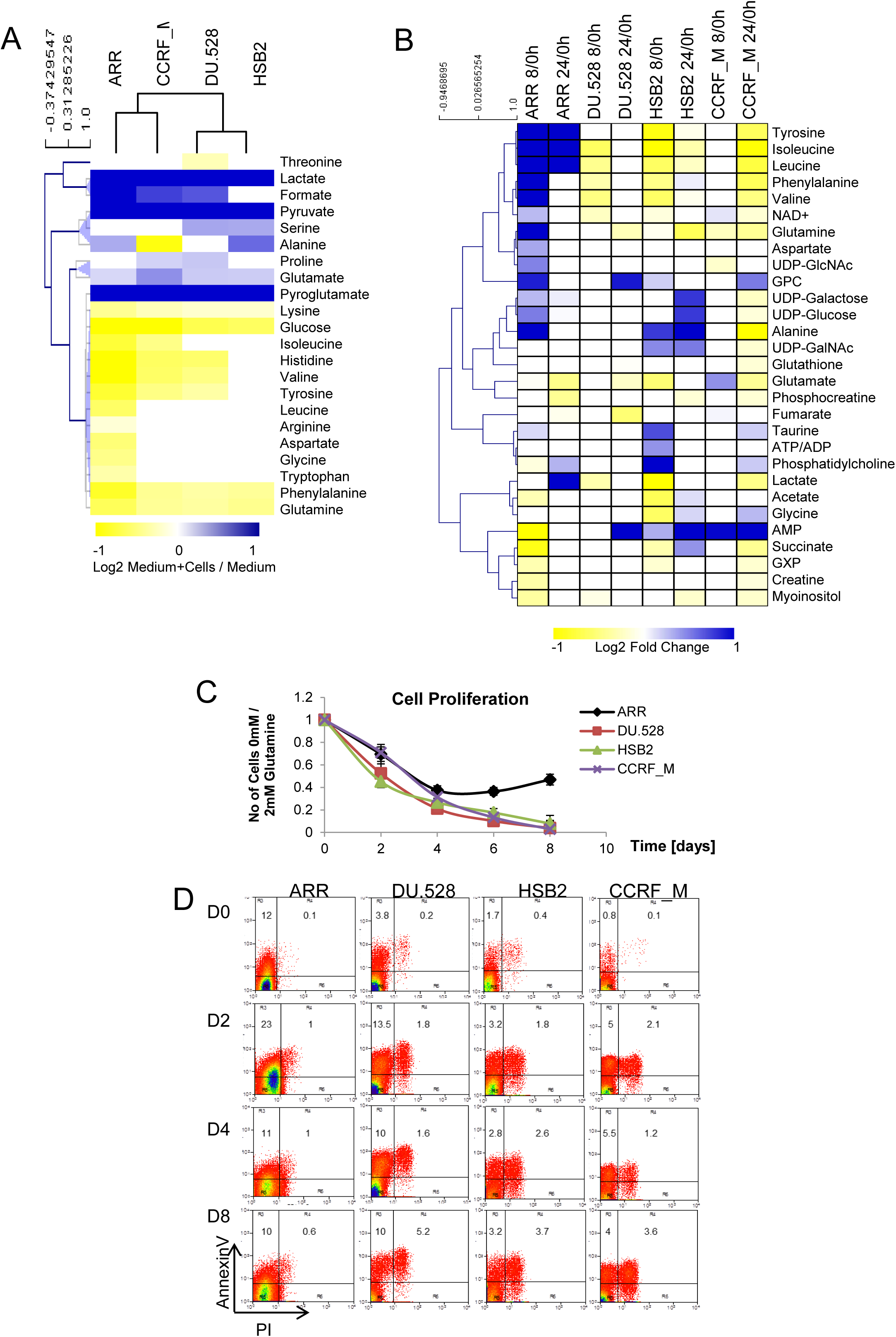
Metabolite levels in T-ALL cells are dynamic and sensitive to mTOR activity. **A)** Heat map showing hierarchical clustering of the growth medium metabolites after 24h. Data are the average of four independent experiments ± StDev. Scale bar represents log2 relative metabolite concentration. Metabolites with significantly different levels are shown. **B)** Abundance of intracellular metabolites at 8h or 24h, relative to the time of medium change (0h). Fold change was presented only for metabolites with significantly different relative levels with p<0.05. GPC, L-Alpha glycerylphosphorylcholine; GalNAc, Acetylgalactosamine, GlcNAc, N-Acetylglucosamine. **C)** Glutamine deprivation inhibits T-ALL cell proliferation. T-ALL cell lines were cultured in medium with 10% FCS, with or without 2mM GlutaMax. Four independent cell cultures were assayed per cell line and each point represents the mean ± StDev. Statistically significant differences were found when ARR was compared to any of the other cell lines at day 6 and 8 (p<0.05). **D)** Glutamine deprivation induces apoptosis in T-ALL cell lines. Scatter plots show flow-cytometry Annexin V and propidium iodide staining by flow-cytometry of cells grown in RPMI media supplemented with 10% FBS in the absence of glutamine. Gating was determined based on the staining of T-ALL cells grown in the RPMI media supplemented with 2mM glutamine. Cells were grown for the indicated number of days (D0-D8) prior to staining and analysis.

To assess how uptake influences intracellular metabolite levels we compared the abundance of intracellular metabolites at the time of growth-media change (0h) to that seen 8h and 24h later (Figure 2B). A similar response to media change in all four cell lines was found only for glutamate and myoinositol, whose intracellular concentration decreased upon media supplementation. Glutamate and myoinositol are important metabolites for every proliferating cell as they are used to fuel cell growth and proliferation. Myoinositol is used as a substrate for membrane synthesis while glutamate feeds into the TCA cycle hence, energy production. Observed glutamine uptake and glutamate utilisation by T-ALL cell lines suggest that glutamine is used to support the cellular demand for glutamate (Figure 2A,B).

### Glutamine is Essential for T-ALL Proliferation and Survival

Next, we wanted to determine the importance of glutamine for T-ALL. We therefore tested the effect of glutamine withdrawal on cellular proliferation. Omitting glutamine from the growth medium, while maintaining all other nutrients, including 10% foetal bovine serum, reduced T-ALL cell numbers by 95% after 8 days of treatment, except for ARR where the proliferation decreased over the first four days, after which ARR resumed proliferation (Figure 2C). Apoptosis assays of the cells grown in glutamine-free medium revealed that the T-ALL cell lines had an increased rate of apoptosis (Figure 2D), showing that glutamine is important for T-ALL cell proliferation and survival.

### Glutamine Fuels Aspartate and Nucleotide Biosynthesis

Glutamine has several significant roles in metabolic processes, such as fuelling the TCA cycle through glutaminolysis, acting as a nitrogen donor in transamination and transamidation reactions, and *de novo* nucleotide synthesis. Aspartate, glycine, glutamate and alanine can be derived from glutamine, and glutamine, glycine and aspartate are used as substrates for nucleotide synthesis. To assess the glutamine contribution to T-ALL metabolic processes we performed tracer experiments using [2,5-^15^N]Glutamine and [3-^13^C]Glutamine. When cells were grown in the presence of [2,5-^15^N]Glutamine, in addition to the expected ^15^N labelled glutamate (data not shown), we observed ^15^N incorporation into the amino group of aspartate and alanine within 8h of treatment for all four cell lines (Figure 3A,S1A). For aspartate, at 0h, a doublet was seen due to coupling to the Hα. At later time points there was increased spectral complexity due to weak coupling of Hβs to ^15^N. The resulting spectrum was a weighted average of a doublet of doublets (from ^15^N-labelled aspartate) and a simple doublet (unlabelled aspartate) (Figure 3A). This demonstrates that glutamine-derived ^15^N was incorporated into aspartate and alanine by transamination of oxaloacetate and pyruvate, respectively.

**Figure 3.**
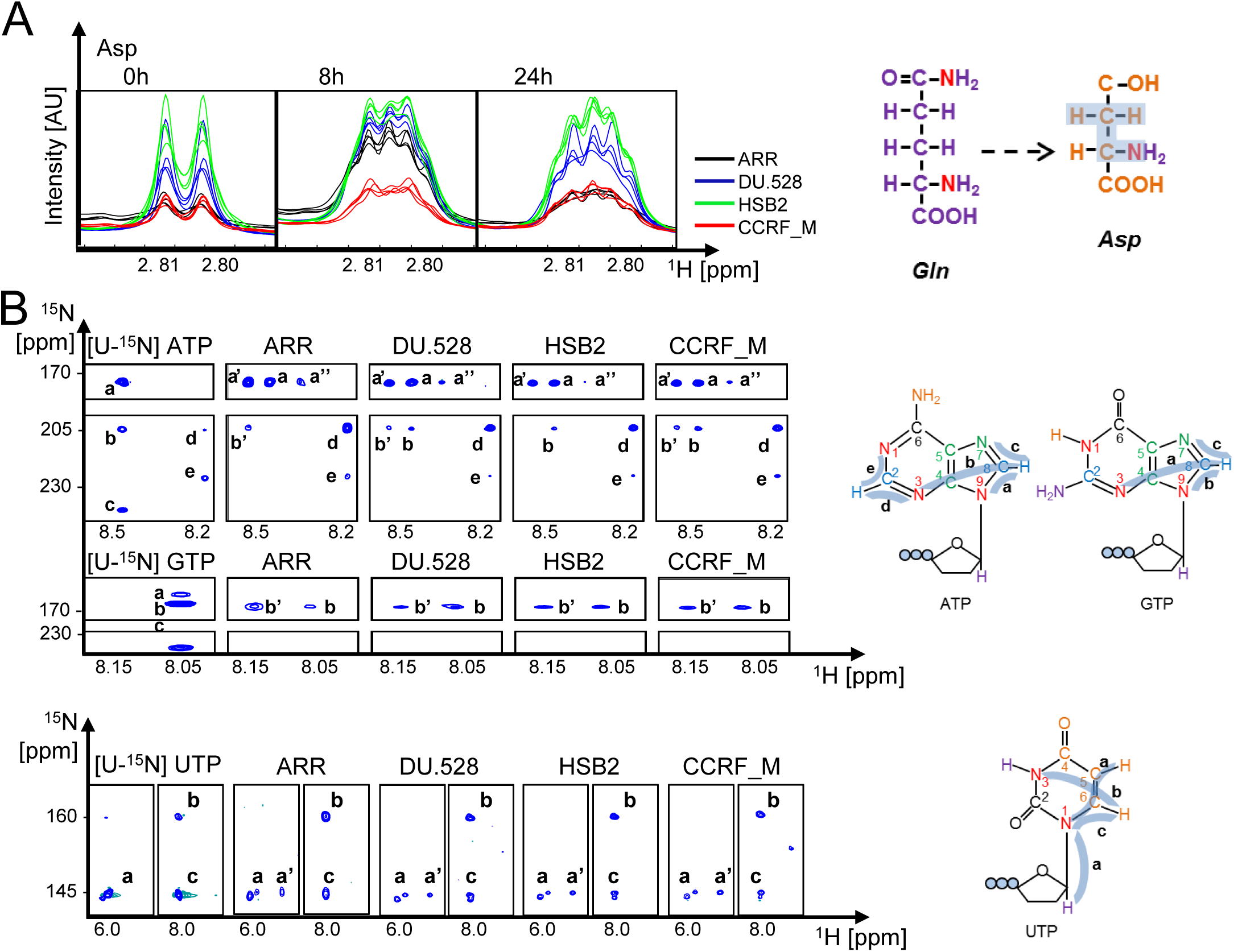
T-ALL cells utilise glutamine-derived nitrogen for *de novo* nucleotide synthesis. Metabolite tracing experiments using T-ALL cell lines that were grown in the presence of 2mM [2,5-^15^N]Glutamine. **A)** Overlay of 1D ^1^H-NMR spectra showing the Hβ-aspartate resonance after 0, 8 and 24h and schematic representations of [2,5-^15^N] Glutamine and the observed [2-^15^N] Aspartate. ^15^N are in red and shading indicates the observed ^3^J scalar couplings between the aspartate Hβs and glutamine-derived ^15^N. The X-axis shows the chemical shift relative to TMSP in ppm and the Y-axis indicates TSA scaled intensity. **B)** Resonances observed in ^1^H-^15^N-HSQC for [U-^15^N]ATP, [U-^15^N]GTP, [U-^15^N UTP standards and for T-ALL cells extracts grown for 24h in the presence of [2,5-^15^N]Glutamine. Resonances are marked by letters **a-e**. Schematics show ATP, GTP and UTP with colour-coded atoms based on the substrate of their origin (glutamine-purple, aspartate-orange, glycine-green, carbonate-black and ^15^N-red). Blue shaded lines indicate observed couplings annotated **a**-**e**.

To assess the contribution of glutamine to nucleotide biosynthesis, we first acquired 2D-^1^H,^15^N heteronuclear single quantum coherence (HSQC) NMR spectra of ^15^N uniformly labelled ATP, GTP and UTP standards, where we observed five ^1^H-^15^N interactions for [U-^15^N]ATP (a-d), three for [U-^15^N]GTP (a-c) and three for [U-^15^N]UTP (a-c) (Figure 3B). Spectra acquired from [2,5-^15^N]Glutamine-labelled T-ALL cells showed ^15^N incorporation into nucleotides at the 1, 3 and 9 positions, the 3 and 9 positions and the 1 and 3 positions for adenine, guanine and uracil respectively (Figure 3B). As expected, transamidation using ^15^N-glutamine resulted in label incorporation at N-3 and N-9 in purines and the N-3 in pyrimidines (Supplementary Illustration (SI)1,2). Additional label incorporation at N-1 in purines and N-3 in pyrimidines is sourced from glutamine-derived ^15^N-aspartate and shows that glutamine ultimately supplies all but one nitrogen in purine and both nitrogens in the pyrimidine ring.

We demonstrated that glutamine transamination of oxaloacetate gives rise to aspartate. It is possible that glutamine supplies oxaloacetate, through glutaminolysis, which would consequently mean that aspartate is completely derived from glutamine. We used [3-^13^C]Glutamine in labelling experiments to test this. Within the TCA cycle, [3-^13^C]Glutamine is converted to [2-^13^C]Fumarate, a symmetrical molecule which is hydrated equally to [2-^13^C]Malate and [3-^13^C]Malate, and further to [2-^13^C] or [3-^13^C]Oxaloacetate and aspartate (SI3). Acquired 2D-^1^H,^13^C HSQC NMR spectra revealed ^13^C incorporation in fumarate, malate, oxaloacetate and aspartate (Figure 4A,S1B and data not shown). Resonances for ^1^H-^13^C moieties were derived from both [2-^13^C] and [3-^13^C] labelling, as shown for aspartate and malate (a and b respectively, Fig 4A,S1B). Quantification of the signals from the labelled samples, relative to the naturally occurring ^13^C in the control samples, showed that within 24h, 40-70% of aspartate was [2-^13^C] or [3-^13^C]Aspartate (Figure 4B). These findings confirm that two molecules of glutamine give rise to a single aspartate molecule; first glutamine supplies the backbone via the TCA cycle, while the second is used for transamination. These metabolic conversions occur in all four tested T-ALL cell lines even though we observed that ARR also uptake aspartate from the media (Figure 2A), suggesting that the aspartate consumption in ARR is not sufficient to support the demand.

**Figure 4.**
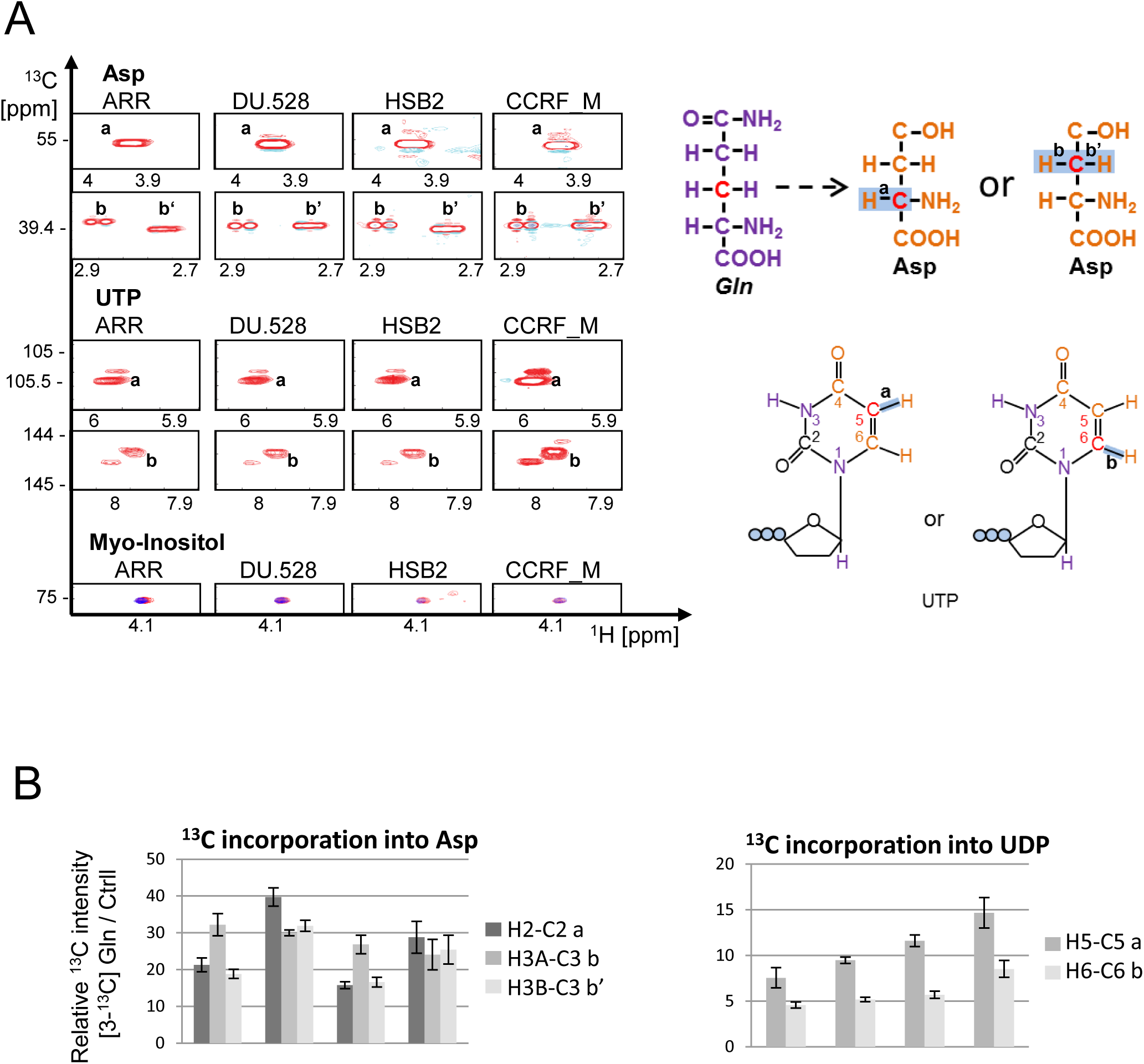
T-ALL cells utilise glutamine-derived carbon for *de novo* nucleotide synthesis. **A)** Resonances observed in ^1^H-^13^C-HSQC for T-ALL cells grown for 24h in the presence of [3-13C]Glutamine. Resonances are marked by letters **a-e**. Schematics on the right show glutamine, aspartate and UTP with colour-coded atoms based on the substrate of their origin. Blue shaded lines indicate observed couplings annotated **a**-**e. B)** Quantification of glutamine-derived aspartate and UDP. Bar graphs show ^13^C signal intensity acquired from the cells grown in the presence of [3-^13^C]Glutamine for 24h, relative to the signal acquired from the naturally occurring ^13^C observed in the extracts from cells grown without the label. Each bar represents one of the interactions shown in the schematics in (A). Data are the average of at least three independent experimental measurements ± StDev.

A further implication of these findings was that carbons derived from glutamine would, via aspartate, get incorporated into pyrimidines, giving rise to [5-^13^C] or [6-^13^C]Uridine (SI2). This was indeed observed in 2D-^1^H,^13^C HSQC NMR spectra (peaks a and b, Figure 4A,B), showing that the carbon at position 4 also originates from glutamine via aspartate. Together, our results show that glutamine serves as a source for all but one of the atoms in the pyrimidine ring. Additionally, examination of the ^13^C incorporation in other detectable metabolites found a significant label accumulation into proline, as would be expected since glutamate is used as a substrate for proline synthesis (Figure S1C).

### Patient-derived T-ALL and T-ALL cell lines have similar metabolic uptake

To determine if glutamine is not only used by patient derived cell lines, but also by primary T-ALL patient isolates, we measured live-cell real-time metabolite uptake by NMR. T-ALL CD34^+^/CD7^+^ cells were isolated from three T-ALL patients at presentation. The flow cytometry profile revealed that samples had cytoplasmic CD3 expression but lacked CD3 surface expression. T-ALL_1 consisted of 60% CD34^+^ blasts, half of which were CD7^+^. T-ALL_2 was predominantly CD7^+^, as well as CD5^+^, CD2^+^, CD38^+^ and CD4^+^, whereas T-ALL_3 was CD7^+^, CD13^+^, CD5^+/-^, CD2^+/-^, CD117^+/-^, while CD1a^-^, CD4^-^ and CD8^-^. CD34^+^ (T-ALL_1) or CD7^+^ (T-ALL_2/3) cells were isolated and 10^6^ cells were resuspended in RPMI medium supplemented with GlutaMAX (L-glutamine/L-alanine dipeptide) and used for measuring metabolite uptake^30^. GlutaMAX is a temperature-stable source of glutamine, which is hydrolysed by peptidases located on the plasma membrane, into L-Glutamine and L-Alanine^31^. Using live-cell NMR we observed for all the patient samples that GlutaMAX concentrations decreased without the equivalent increase in glutamine availability showing that patient-derived T-ALL utilise glutamine (Figure 5A).

**Figure 5.**
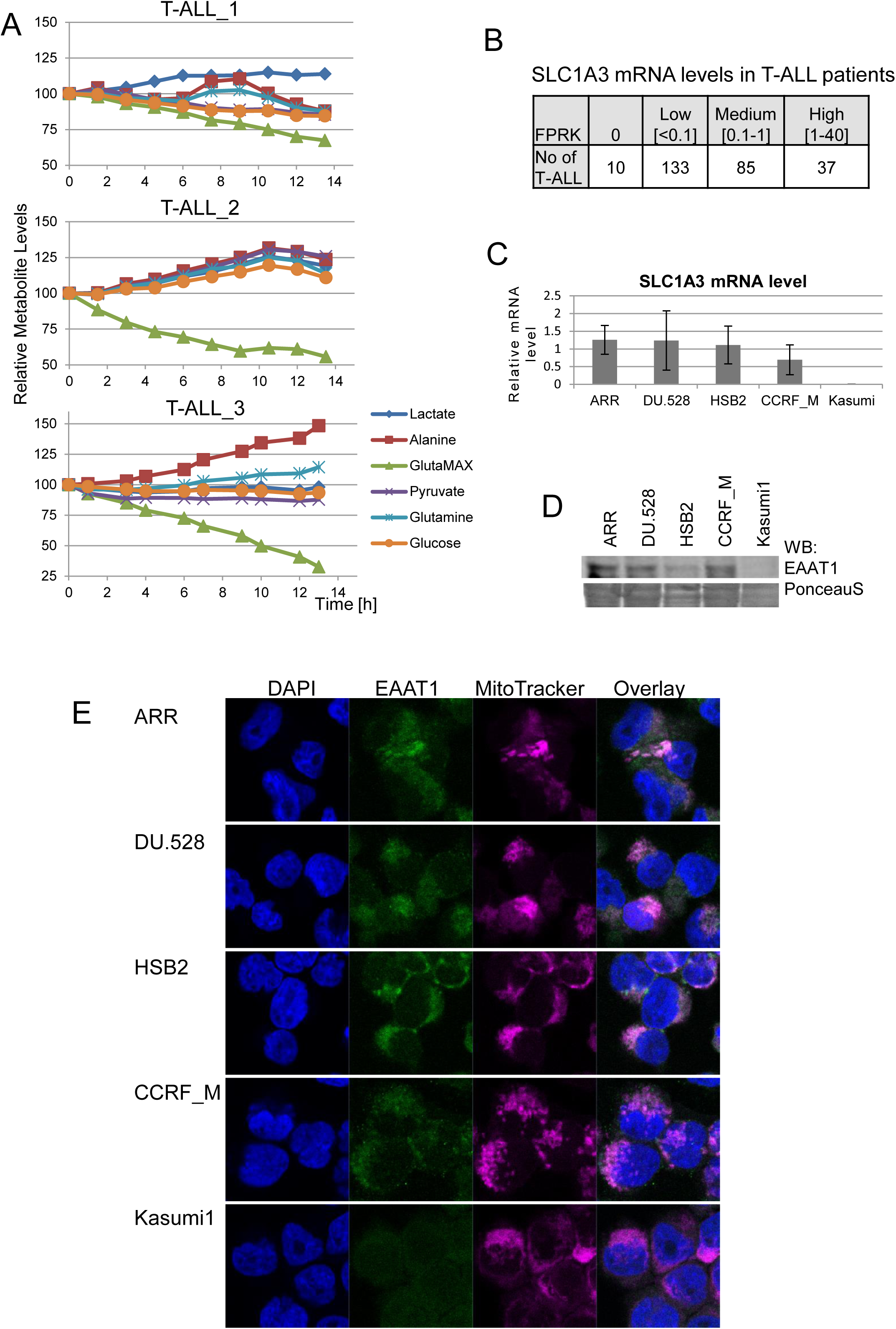
EAAT1 is expressed in T-ALL and localised in mitochondria. **A)** Patient derived T-ALL cells have similar metabolic uptake to T-ALL cell lines. Only metabolites with changing concentration are illustrated. **B)** Summary of the *SLC1A3* expression levels as assessed by FPKM values from the RNA-seq of 265 patient samples. Data are grouped by the level of expression. **C)** *SLC1A3* mRNA expression level in T-ALL cell lines and AML cell line Kasumi-1, relative to rRNA level assessed by qPCR. **D)** Western blot analysis of EAAT1 protein level using 150 µg total cell extract. PonceauS shows equal loading. **E)** Immuno-fluorescent imaging shows that EAAT1 (green) co-localises with MitoTracker Red CMXRos (magenta) in T-ALL but not Kasumi-1.

### T-ALL cells express EAAT1 in the mitochondria

Our finding that T-ALL rely on deriving their aspartate from glutamine instead of sourcing it from the media, highlights that this metabolic step is a T-ALL liability. Therefore, we focused on identifying potential therapeutic targets for T-ALL treatment within the metabolic processes supporting glutamine-derived aspartate synthesis (SI3). Inhibiting the glutamate-aspartate anti-port across the mitochondrial membrane would be a good strategy for targeting T-ALL as it is not directed towards the enzymatic conversions that would restrict aspartate’s availability for nucleotide synthesis in healthy cells. Aspartate export and glutamate import into mitochondria is known to be facilitated by 3 different antiporters *SLC1A3, SLC25A12* and *SLC25A13* ^32,33^. While *SLC25A12* and *SLC25A13* are expressed in most tissues, the expression of *SLC1A3* gene is mainly restricted to the CNS^14,34^. Analysis of our RNAseq data, as well as published gene expression data of 265 T-ALL patient samples, revealed that *SLC1A3* is expressed in our model cell lines and in 95% of patient samples (Figure 5B,C)^2^. Additionally, we observed that EAAT1, the protein encoded by *SLC1A3*, was detected in the mitochondria of the T-ALL cell lines but not in the acute myeloid leukaemia (AML) cell line Kasumi-1, which does not express SLC1A3 (Figure 5C,D,E)^35^. Together, these results show that T-ALL cells express the high affinity glutamate-aspartate antiporter EAAT1.

### SLC1A3/EAAT1 is essential for oncogenic de novo nucleotide synthesis

To assess the importance of EAAT1 for T-ALL survival we performed SLC1A3 knock-down using shRNA. Five shSLC1A3 were designed and tested for their capacity to supress the expression of SLC1A3 cDNA. Mouse fibroblasts were co-transduced with retrovirus expressing SLC1A3-IRES-GFP and shRNAs. shSLC1A3 efficiency was measured relative to the negative control, shFF3, and the positive control shGFP. shSLC1A3_1 and shSLC1A3_2 had the capacity to supress EAAT1 protein levels similar to shGFP, while the negative control shFF3 did not have any effect on EAAT1 protein levels (Figure 6A). shSLC1A3_1 and shSLC1A3_2 were cloned into a PiggyBac backbone that supports stable doxycycline-inducible shRNA expression. Induction of shSLC1A3 led to rapid cell death within 5 days from the start of the doxycycline treatment (Figure 6B).

**Figure 6.**
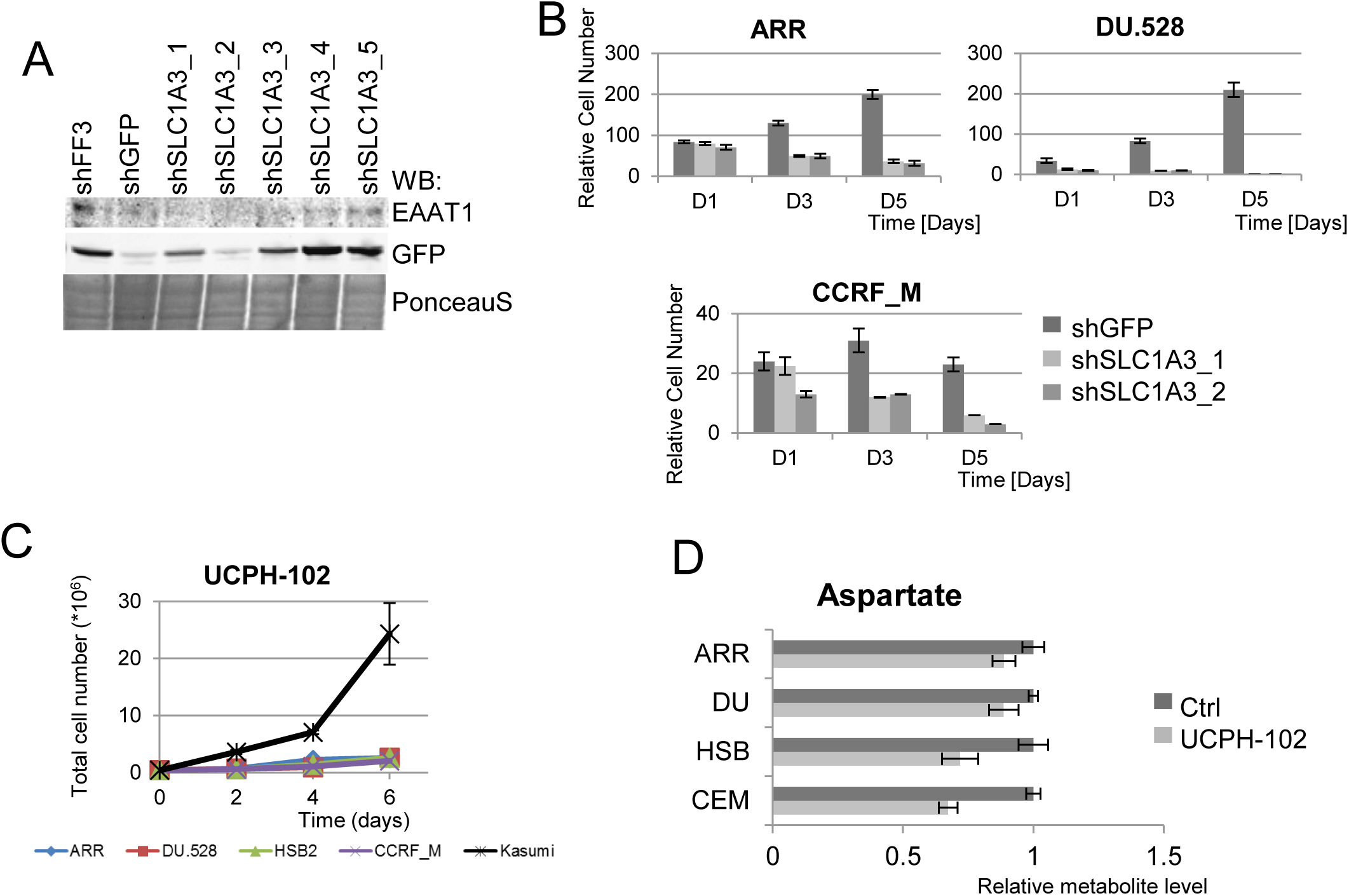
EAAT1 is essential for T-ALL proliferation and survival. **A)** Protein level of EAAT1 and the reporter GFP protein upon the suppression of SLC1A3-IRES-GFP mRNA by shFF3 (negative control), shGFP (positive control) and five different shSLC1A3_1-5. Experiment was performed in duplicate. PonceauS staining illustrates equal loading. **B)** Knock-down of the SLC1A3 gene by shSLC1A3_1 and shSLC1A3_2 leads to ARR, DU.528 and CCRF_M cell death. **C)** To test the effect of EAAT1 inhibition, T-ALL and AML cells were cultured for six days in the presence of vehicle (DMSO) or 25 µM UCPH-102. Each data point is an average of three independent measurements ± StDev. **D)** UCPH-102-induced changes in aspartate uptake. T-ALL cell lines were grown in the presence of 25µM UCPH-102 or vehicle (DMSO) for 48h. Relative aspartate levels are the mean of four independent experiments. Two-tailed t-test identified the difference in aspartate concentration between the cells grown with vehicle and UCPH-102 as statistically significant (p<0.05) in all four cell lines.

With the purpose to inhibit EAAT1 function in the CNS, the allosteric inhibitor UCPH-102 was developed^36,37^. Treatment with UCPH-102 had an anti-proliferative effect on T-ALL cells but not on the EAAT1-negative AML cell line, showing that EAAT1 function is required for T-ALL proliferation (Figure 6C). Next we measured the effect of UCPH-102-mediated EAAT1 inhibition on metabolite uptake/release and found that UCPH-102 treatment significantly increased aspartate consumption in all four T-ALL cell lines (Figure 6D), implying that T-ALL have the capacity to uptake aspartate but that this is not sufficient to compensate for the loss of EAAT1 activity. Altogether, our results show that oncogenic nucleotide production depends on glutamine uptake and that extracellular aspartate cannot support T-ALL proliferation and survival.

### EAAT1 is required for T-ALL xenograft development

To test the importance of EAAT1 during disease progression in a mouse xenograft model, CRRF-CEM cells, carrying doxycycline-inducible shSLC1A3_2 or shGFP in conjunction with TdTomato, were injected into the tail vein of NSG mice. When engraftment was apparent at approximately 1% of total CD45^+^ blood mononuclear cells, the diet was supplemented with doxyxycline to induce shRNA expression. After 6 days, significantly less human CD45^+^ cells were detected in mice with shSLC1A3-expressing CCRF-CEM cells, compared to mice that received shGFP-expressing cells (Figure 7A). Doxycycline treatment resulted in a survival advantage for the shSLC1A3 mice compared to the shGFP control mice (Figure 7B). While all human CD45^+^ shGFP-expressing cells were TdTomato-positive, this was the case for only 40% of human CD45^+^ cells in mice with shSLC1A3-expressing cells, indicating the outgrow of cells that did no longer express the shSLC1A3 construct (Figure 7C). These *in vivo* results show that EAAT1 gives a proliferative advantage to T-ALL and validates EAAT1 as therapeutic target.

**Figure 7.**
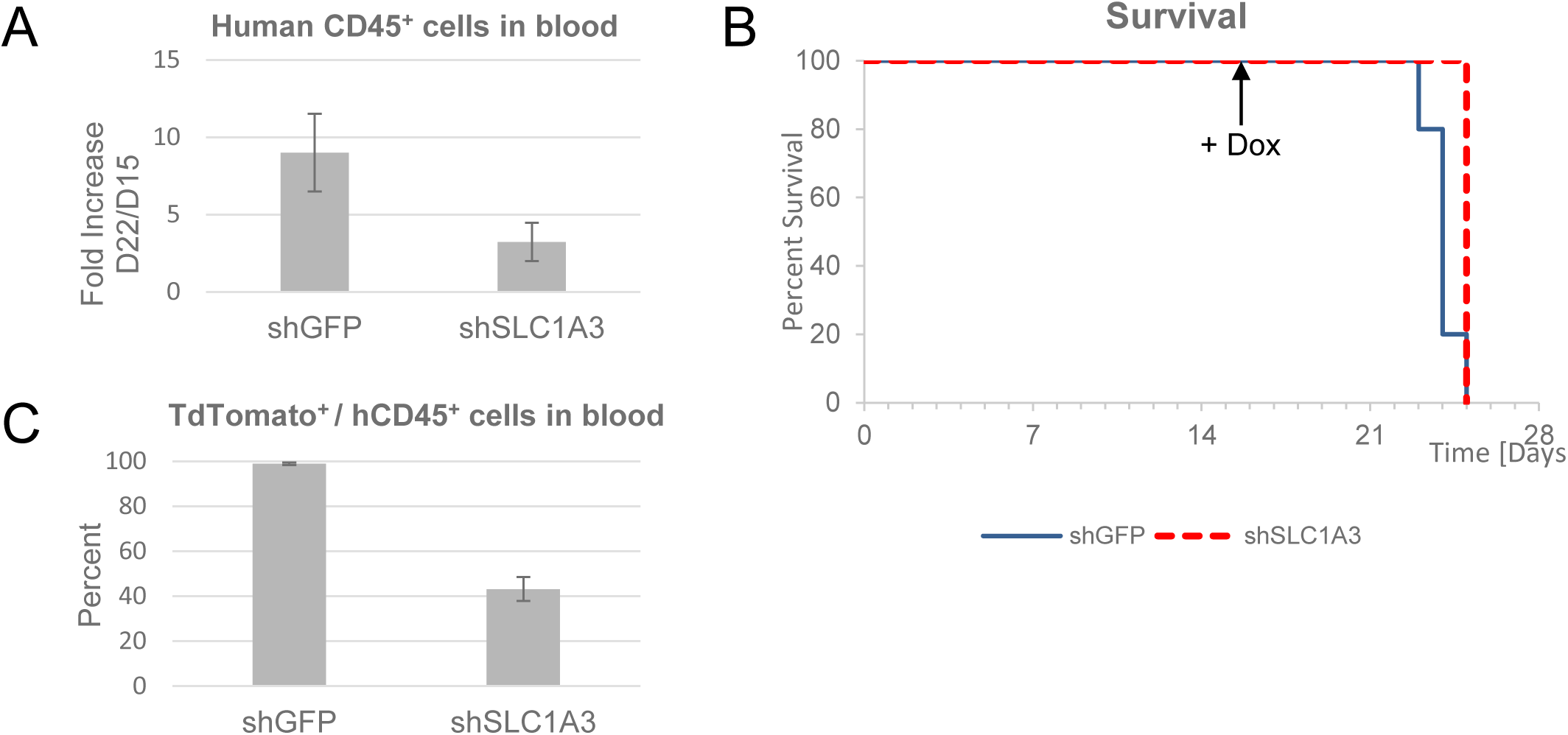
EAAT1 is required for T-ALL xenograft development. **A)** Mice were injected (iv) with 3 ×10^5^ cells CCRF-CEM cells carrying doxycycline-inducible shSLC1A3_2 or shGFP. At day 16, the food was supplemented with doxycycline. The bar graph shows the number of human CD45^+^ cells in the two cohorts of mice (n=5) after 6 days of doxycycline treatment. Two-tailed t-test identified the difference between the two cohorts as significantly different (p<0.05). **B)** Kaplan Meier curve comparing the survival of mice injected with CCRF-CEM cells carrying doxycycline-inducible shSLC1A3 or shGFP. The induction of shRNA expression through addition of doxycycline in the food on day 16 is indicated. **C)** The number of TdTomato-positive cells relative to the total number of hCD45^+^ cells at day 6.

## Discussion

Understanding the metabolic pathways that support oncogenic proliferation can help to identify cancer-specific processes and rate-limiting steps that can be used for developing new therapies. We show that in T-ALL cells, glutamine is converted to glutamate that enters the mitochondria. Here, glutamate dehydrogenase (GLUD1,2) uses it to generate α-ketoglutarate that will enter the TCA cycle (Figure 8). Mitochondrial glutamate oxaloacetate transaminase (GOT2) uses glutamate as a donor for the amino group, which is transferred to the TCA cycle intermediate oxaloacetate, resulting in the generation of aspartate. Aspartate is then transported out of the mitochondria in exchange for a new glutamate molecule. Together, glutamine and aspartate are used as substrates for nucleotide synthesis (Figure 8).

**Figure 8.**
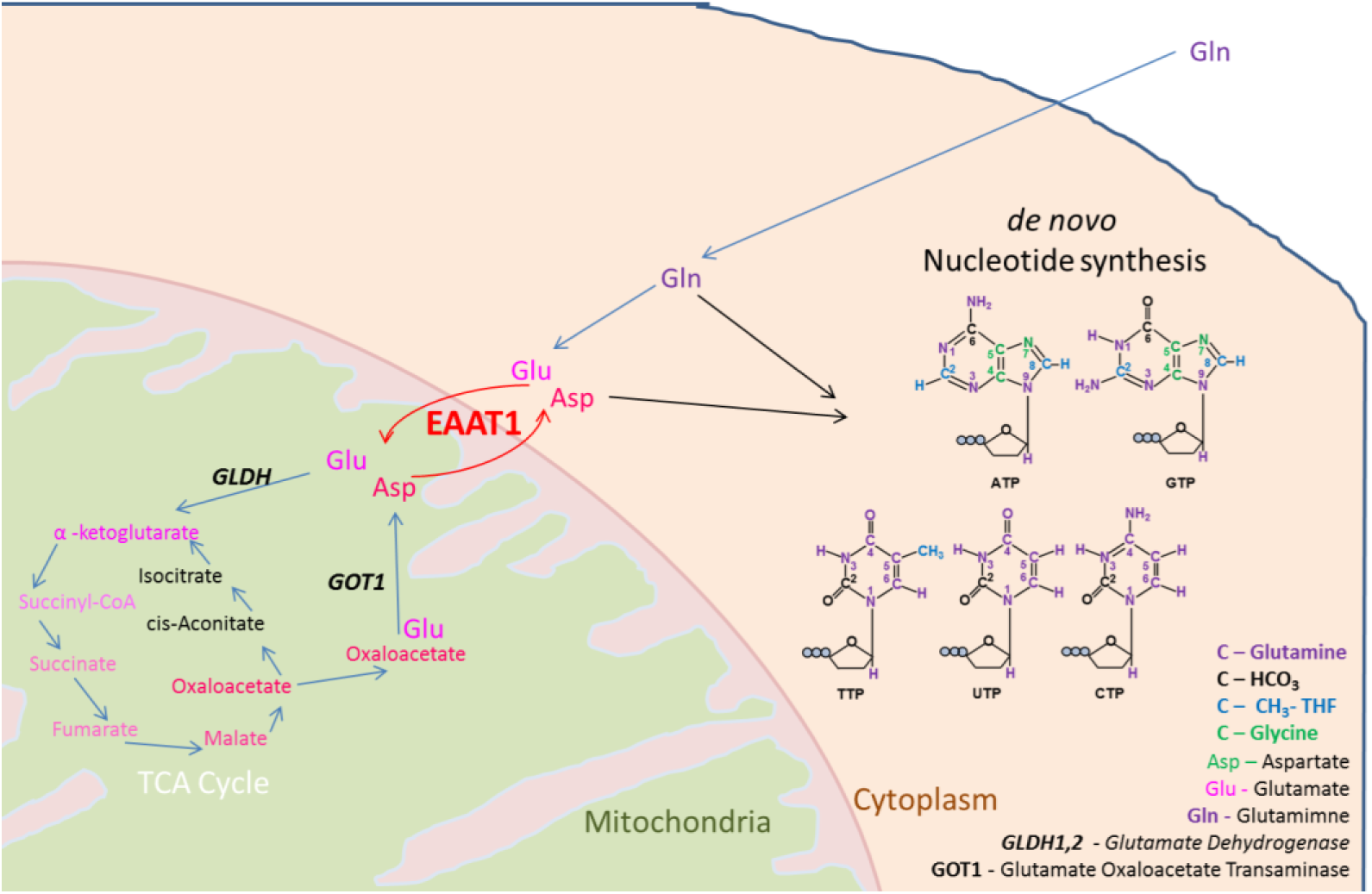
Model illustrating the function of mitochondrial EAAT1.

Similar processes are likely to occur in other cancers. Indeed, a recent study of genetic cancer dependencies has shown that T-ALL and several other cancer types are dependent on SLC1A3 expression^38^. Additionally, EAAT1 has been implicated in supporting proliferation in several solid cancer cell lines^39,40^. These studies showed a role for EAAT1 in the uptake of aspartate from the medium, especially under conditions of glutamine deprivation or asparaginase treatment. In both studies, removing EAAT1 under normal cell culture conditions showed little or no effect, in contrast to our work with T-ALL cells. This difference can be explained as the solid tumour cell lines were mainly dependent on EAAT1 for aspartate/glutamate uptake from the environment and the lack thereof resulted in a combinatorial effect on the TCA cycle, the electron transport chain, and *de novo* glutamine/glutamate and nucleotide synthesis. T-ALL cells on the other hand are dependent on the availability of glutamine. They rely on the function of EAAT1 at the mitochondria, where the glutamate/aspartate antiport is required for *de novo* nucleotide production. Our isotope tracing experiments show that the carbon and nitrogen atoms of glutamine are used for aspartate, purine and pyrimidine biosynthesis. Sun *et al*. also showed that asparaginase treatment resulted in a cellular depletion of glutamine/glutamate, possibly explaining why asparaginase treatment is often effective in treatment of T-ALL^41^.

EAAT1 is normally present on the plasma membrane of neurons and glia in the CNS, where it uptakes glutamate from the glutamatergic synapses^13^. Based on the RNA expression and protein localisation assessed with three different specific antibodies, the Human Protein Atlas Database reports that EAAT1 is not found at significant levels outside of the CNS (https://www.proteinatlas.org/ENSG00000079215-SLC1A3/tissue)^42^. However, EAAT1 expression has been reported in neonatal cardiomyocytes where, similar to T-ALL, it is also localised at the mitochondria^32^.

Altogether, we show that SLC1A3 is aberrantly expressed in T-ALL cells, where it supports *de novo* nucleotide production which uses glutamine as the main substrate, and that EAAT1 is required for T-ALL proliferation, identifying it as a relevant therapeutic target. Unfortunately, the selective and potent EAAT1 inhibitor UCPH-102, which has been developed for the inhibition of EAAT1 in the CNS, is not suitable for *in vivo* use and therefore new therapeutic agents need to be developed that can target the oncogenic function of EAAT1.

## Supporting information

Supplementary Illustrations

## Acknowledgments

We would like to thank Dr. A.W. Langerak, (Erasmus Medical Centre, Rotterdam, NL) for the provision of ARR and DU.528 cell lines and Prof P.N. Cockerill (University of Birmingham, UK) for HSB2 and CCRF-CEM. We would like to thank Dr. M. McGrew (University of Edinburgh, UK) for the PB_tet-on_Apple_shGFP plasmid and Prof. L. Bunch for pcDNA3-EAAT1 plasmid. We would like to acknowledge the University of Birmingham FACS facility for cell sorting and the Biomolecular NMR Facility at the Henry Wellcome Building for Nuclear Magnetic Resonance (HWB-NMR), University of Birmingham. This work was supported by Bloodwise, through a Bennett Fellowship to M.H. [11002], the Medical Research Council and the University of Birmingham.

## Authorship Contributions

Project Planning, Experimental Design, V.S.S., M.H.; NMR data acquisition and analysis V.S.S., M.A.C.R., J.R., U.G., C.L.; Animal experiments, V.S.S., T.A.P., M.H.; Other Experimentation, Data Acquisition, Processing and Analyses V.S.S., S.AO., M.H.; Statistical analysis V.S.S.; Provision of patient samples S.P., G.P.; Reagents provision S.S.; Figures V.S.S., M.H.; Writing of the Manuscript, V.S.S., M.A.C.R., M.H.

The authors declare no conflicts of interests.

