## Supplementary figures and images for "T-ALL Proliferation Relies on Glutamine Uptake and EAAT1-dependent Conversion of Glutamine to Aspartate and Nucleotides"

### Supplementary figures

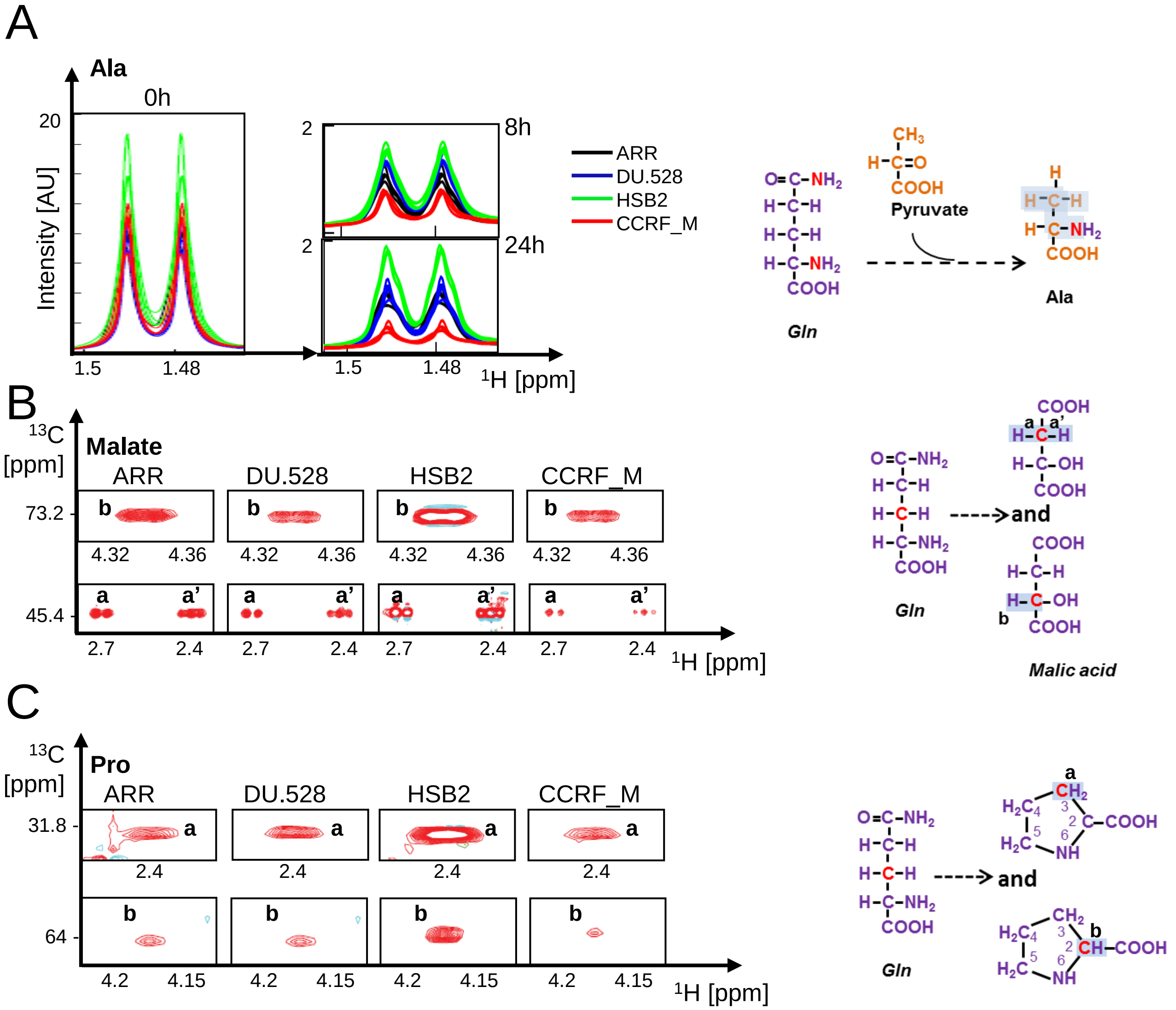

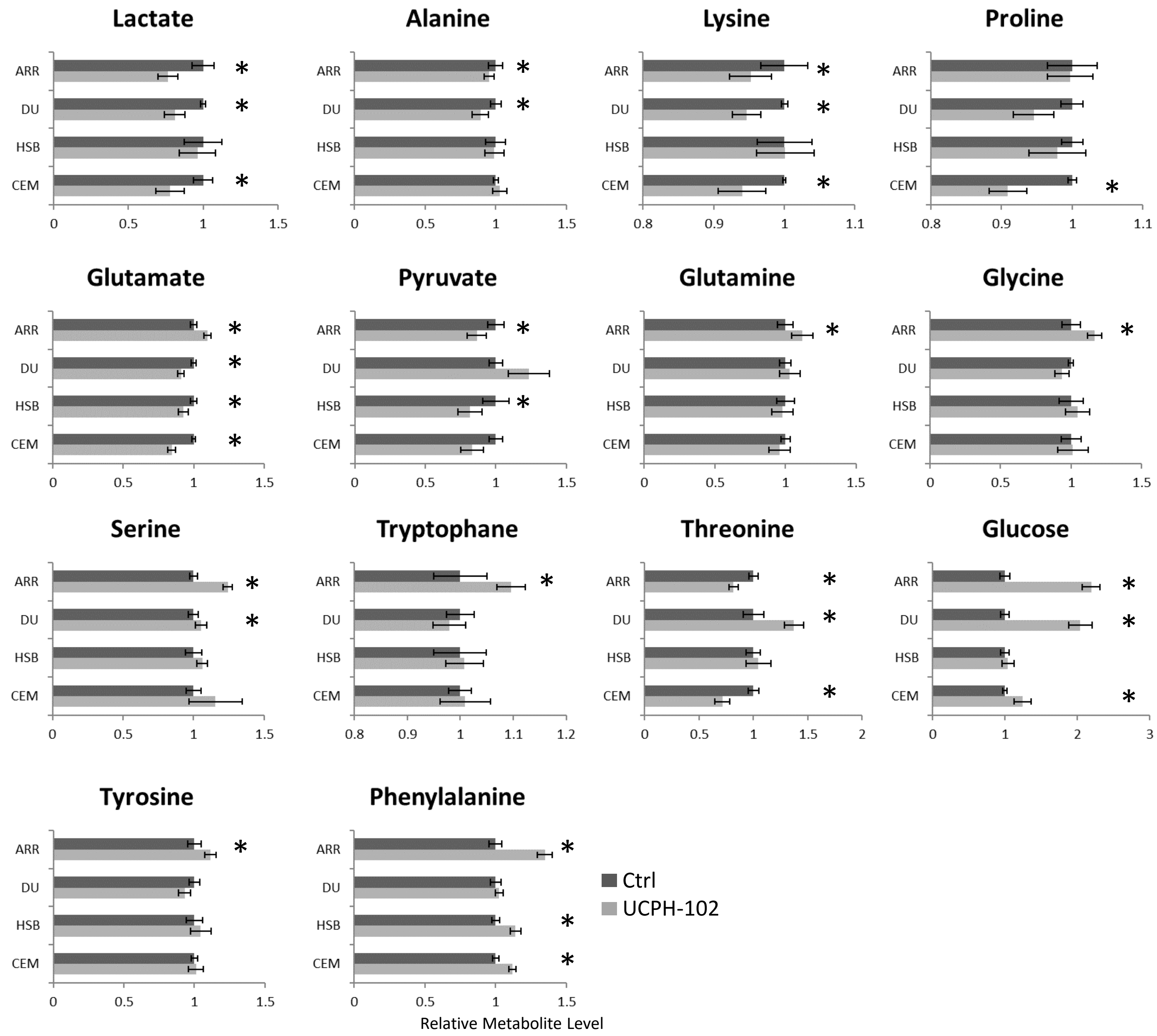
