## Supplementary figure legends for "T-ALL Proliferation Relies on Glutamine Uptake and EAAT1-dependent Conversion of Glutamine to Aspartate and Nucleotides"

### Supplemental Figure Legends

**Figure S1.** Metabolite tracing experiments using [2,5-$^{15}$N]Glutamine and [3-$^{13}$C]Glutamine. T-ALL cell lines were grown in the presence of 2mM [2,5-$^{15}$N]Glutamine) (A) or [3-$^{13}$C]Glutamine) (B-D). **A)** Overlay of 1D $^{1}$H-NMR spectra showing the Hβ-alanine resonance after 0, 8 and 24h and schematic representations of [2,5-$^{15}$N]Glutamine and the observed [2-$^{15}$N]Alanine. $^{15}$N are in red and shading indicates the observed $^{3}$J scalar couplings between the alanine Hβs and glutamine-derived $^{15}$N. The X-axis shows the chemical shift relative to TMSP in ppm and the Y-axis indicates TSA scaled intensity. **B,C)** Resonances observed in $^{1}$H-$^{13}$C-HSQC for T-ALL cells grown in the presence of [3-13C]Glutamine for 24h. Resonances are marked by letters **a-b**. Schematics on the right show malate, proline and arginine with colour-coded atoms based on the substrate of their origin. Blue shaded lines indicate observed couplings.

**Figure S2.** Changes in metabolite uptake and release during EAAT1 inhibition. Bar graph showing metabolite levels from the T-ALL cells cultured for 48h in the presence of 25µM UCPH102 or vehicle (DMSO) from four independent samples and relative to the media control. Data are the average ± StDev. Metabolites with significantly different levels between the UCPH102 and the control are indicated with (*). Significance was based on a two-tailed t-test with $p<0.05$.
