## Supplementary Illustrations for "T-ALL Proliferation Relies on Glutamine Uptake and EAAT1-dependent Conversion of Glutamine to Aspartate and Nucleotides"

Supplementary Illustration SI1

### Purine *De Novo* Synthesis

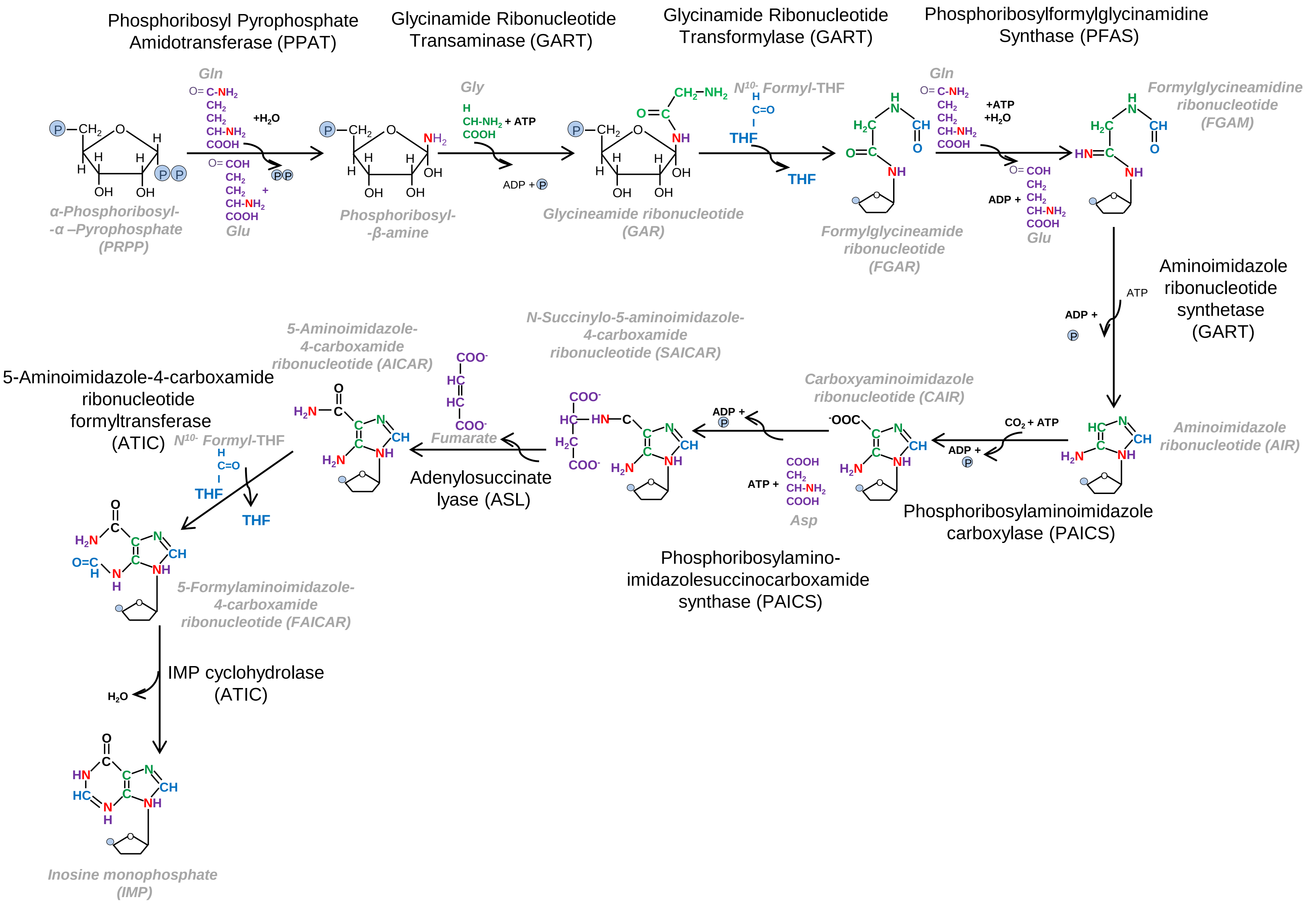

Supplementary Illustration SI2

### Pyrimidine *De Novo* Synthesis

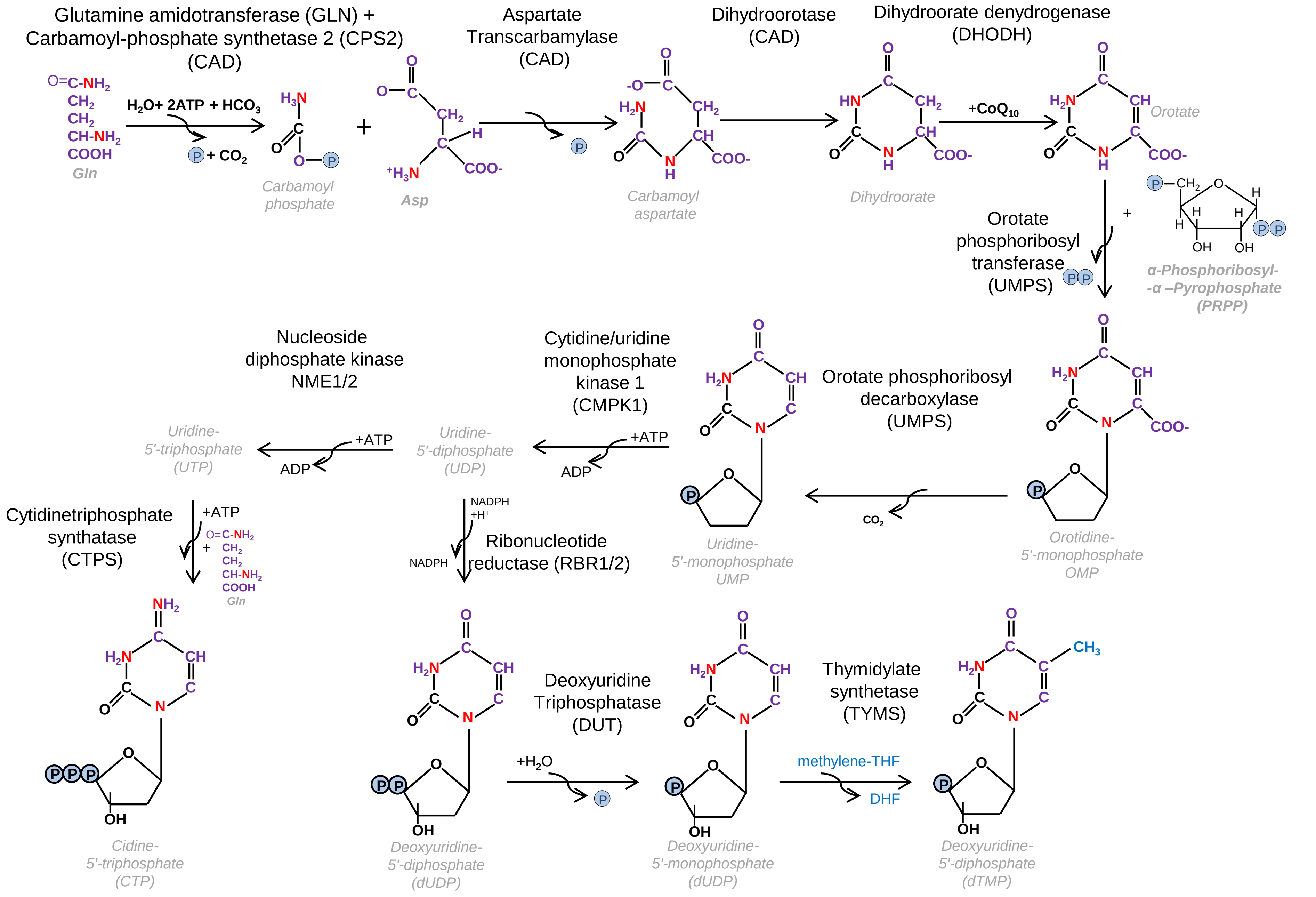

Supplementary Illustration SI3

### *Glutamine to Aspartate*

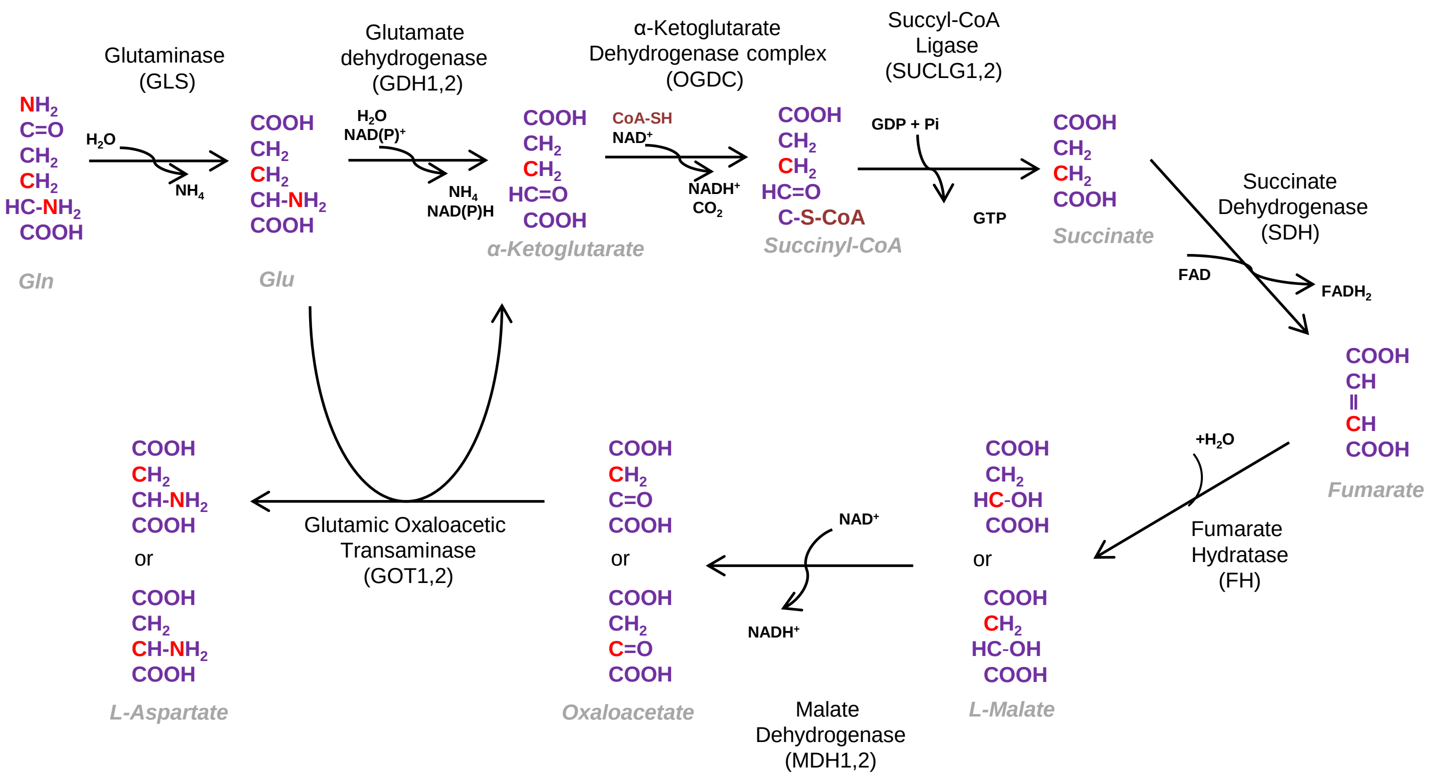
